# Circular RNA-associated QTLs show stronger association with splicing-QTLs than with expression-QTLs

**DOI:** 10.64898/2026.05.29.728707

**Authors:** Aitor Zabala, Alex M Ascensión, Saioa Gs Iñiguez, Leire Iparraguirre, Eduardo Andrés-León, Fuencisla Matesanz, David Otaegui, Maider Muñoz-Culla

## Abstract

**Introduction:** Circular RNA quantitative trait locus (circQTLs) have emerged as a class of regulatory variants, but their mechanistic basis remains poorly characterized. Understanding how genetic variation influences circRNA biogenesis is essential to clarify their role in post-transcriptional gene regulation.

**Methods:** We systematically compared circQTLs with matched splicing (sQTL) and expression (eQTL) datasets. Using bootstrap-based Jaccard similarity analyses, we quantified genomic overlap patterns and assessed their statistical significance. We further validated these findings across independent circQTL studies. In addition, we analyzed the genomic distribution of circQTLs to identify enrichment patterns across functional genomic regions.

**Results:** circQTLs exhibited a statistically significant but modestly stronger genomic overlap with sQTLs compared to eQTLs. This pattern was consistent across independent datasets despite limited reproducibility of individual circQTL signals. Genomic annotation revealed distinct distributional patterns, including depletion in exonic regions and relative enrichment in non-coding genomic contexts compared to other QTL classes.

**Discussion:** Together, these results suggest that circRNA-associated regulatory variation is preferentially linked to splicing-related mechanisms rather than transcriptional control of host genes. However, the modest effect size indicates that this relationship is not exclusive, and likely reflects a mixture of shared splice-site regulatory effects and additional mechanisms specific to back-splicing that are not captured by conventional sQTL or eQTL frameworks. This dual architecture positions circRNA biogenesis at the interface between splicing dynamics, RNA structure, and higher-order genomic organization, supporting circQTLs as a distinct layer of post-transcriptional gene regulation.

## Introduction

Circular RNAs (circRNAs) constitute a conserved and large class of RNA molecules characterized by a covalently closed structure generated through back-splicing, a non-canonical splicing reaction in which a downstream splice donor is ligated to an upstream splice acceptor forming their characteristic back-spliced junction (BSJ) (Wang and Wang, 2015; Chen and Yang, 2015). Unlike linear messenger RNA (mRNA), circRNAs lack free 5’ and 3’ ends, rendering them resistant to exonuclease-mediated degradation and conferring remarkable stability (Jeck et al., 2013). circRNAs have been implicated in diverse regulatory functions, including interaction with micro RNA (miRNA) and RNA-binding protein (RBP), modulation of transcription, and translational capacity (Hansen et al., 2013; Memczak et al., 2013; Jeck et al., 2013). Their dysregulation has been associated with neurological disorders, cancer, and immune-mediated diseases (Iparraguirre et al., 2017; Meng et al., 2017; Iparraguirre et al., 2020, 2021), highlighting the need to understand the regulatory mechanisms behind circRNA expression.

Back-splicing occurs co-transcriptionally and competes directly with canonical precursor messenger RNA (pre-mRNA) splicing for the same splice sites and spliceosomal machinery. Unlike canonical splicing, which proceeds through stepwise exon definition followed by a transition to intron definition and spliceosomal activation (Sharma et al., 2008; De Conti et al., 2013; Fox-Walsh et al., 2005), back-splicing requires spliceosome assembly across exons and therefore does not necessarily undergo this canonical transition, or does so in an alternative configuration, requiring distinct regulatory mechanisms (Wang and Wang, 2015). The efficiency of circularization depends on multiple factors, including splice-site strength, intronic architecture, RNA secondary structure, transcriptional elongation kinetics, and the activity of specific RNA-binding proteins. As a result, the same pre-mRNA molecule can give rise to either linear or circular RNA isoforms, establishing a regulated balance between protein-coding and non-coding outputs from a single genomic locus (Zhang et al., 2016; Liang et al., 2017).

Genome-wide association studies (GWASs) have identified thousands of genetic variants associated with human complex traits (Abdellaoui et al., 2023). Most are in non-coding regions, suggesting effects primarily through regulatory mechanisms that shape gene expression and RNA processing (Edwards et al., 2013; Pickrell et al., 2010). Expression quantitative trait locus (eQTLs) have been extensively mapped across human tissues and predominantly localize to promoters, enhancers, and other regulatory elements that influence transcriptional output (Consortium, 2020). In parallel, splicing quantitative trait locus (sQTLs) capture genetic variants that alter splicing decisions, such as exon inclusion, splice-site usage, or intron retention, and are enriched within exons and intronic regions near splice junctions (Li et al., 2016; Takata et al., 2017; Garrido-Martín et al., 2021). Large-scale resources such as the Genotype-Tissue Expression Portal (GTEx) have established eQTLs and sQTLs as distinct yet complementary layers of genetic regulation (Consortium, 2020). Recent brain transcriptomics studies have identified circRNA-eQTLs pairs enriched at canonical splice sites, revealing their independent colocalization with GWAS signals for complex traits (Chen et al., 2026), motivating further studying of backsplicing regulation.

Recently, circular RNA quantitative trait locus (circQTLs), defined as SNPs associated with changes in the expression of nearby circRNAs, have emerged as a new class of functional variants, although this field remains nascent. Initial studies have identified thousands of circQTLs across brain, blood, and other tissues, and several studies have linked them to complex diseases, including neurodevelopmental and neuropsychiatric disorders as well as metabolic and inflammatory conditions (Ahmed et al., 2019; Liu et al., 2019; Mai et al., 2022; Iñiguez et al., 2026),

Beyond their regulatory relevance, circQTLs are increasingly implicated in disease biology, where genetic regulation of circRNA abundance may contribute to pathogenic processes independently of canonical gene expression effects (Iñiguez et al., 2026). Notably, only a subset of circQTLs also influence mRNA expression (Ahmed et al., 2019), suggesting that many circQTLs act specifically at the level of circRNA formation rather than general transcript abundance. However, the regulatory basis of circQTLs remains poorly characterized. In particular, most circQTL studies have not systematically examined their genomic colocalization with well-established eQTL and sQTL maps. This limits our understanding of whether variation in circRNA expression is primarily driven by transcriptional regulation or by splicing-mediated mechanisms. Given that circRNA biogenesis depends on back-splicing, which competes with canonical splicing for shared splice sites and regulatory machinery, disentangling these contributions is essential to clarify how genetic variation shapes circRNA production.

Given the mechanistic dependence of circRNA biogenesis on splicing, we hypothesized that circQTLs would exhibit stronger genomic colocalization with sQTLs than eQTLs. Variants affecting splice-site strength, intronic pairing, or spliceosomal efficiency could shift the balance between linear splicing and circularization, producing coordinated genetic effects on circRNA abundance and splicing phenotypes. At the same time, back-splicing is regulated by mechanisms that differ from those governing canonical splicing, raising the possibility that a subset of circQTLs operates independently of detectable changes in canonical splicing or overall gene expression.

In this study, we systematically investigate the relationship between circQTLs, sQTLs, and eQTLs by mapping circQTLs in peripheral blood leukocytes and quantifying their genomic overlap with matched splicing and expression QTL datasets. Using bootstrap-based Jaccard similarity analyses and external validation across independent circQTL studies, we test whether circQTLs are more closely associated with splicing-related regulatory variation than with transcriptional control. In addition, we examine the genomic distribution of circQTLs to identify features that distinguish them from other QTL types. Together, this study aims to clarify the regulatory architecture of circRNA expression and to position circQTLs within the broader framework of genetic regulation of RNA processing.

## Results

### circQTLs show a stronger overlap with sQTLs than with eQTLs

CircRNA biogenesis depends on back-splicing, a process mechanistically linked to pre-mRNA splicing and influenced by transcriptional regulation. Consequently, genetic variants associated with circRNA expression may act through splicing-related mechanisms, transcriptional control, or a combination of both. Determining whether circQTLs preferentially overlap with sQTLs or eQTLs is therefore essential to elucidate the regulatory basis of circRNA expression.

To investigate this relationship, we examined the genomic overlap between circQTLs, sQTLs, and eQTLs. Among 38,477 circQTLs, 377,632 sQTLs, and 1,277,232 eQTLs, we found that 10.43% of circQTLs overlapped with sQTLs, 32.49% overlapped with eQTLs, and 8.63% overlapped with both classes (**Figure 1A**). However, the large differences in dataset size complicate direct comparisons of the relative association of circQTLs with sQTLs versus eQTLs. To address this limitation, we applied a bootstrap resampling strategy, generating random samples of 20,000 variants from each QTL type per iteration over 1,000 iterations (described in **Materials and Methods**). Comparison of Jaccard indices revealed a statistically significant but modestly stronger overlap between circQTLs and sQTLs (median = 2.373 *×* 10^−3^) than between circQTLs and eQTLs (median = 2.207 *×* 10^−3^; Mann–Whitney U test: *U* = 6.74 *×* 10^5^, *p* = 3.176 *×* 10^−41^, *r* = 0.301; **Figure 1B, Table S1**, and **Table S2**).

**Figure 1.**
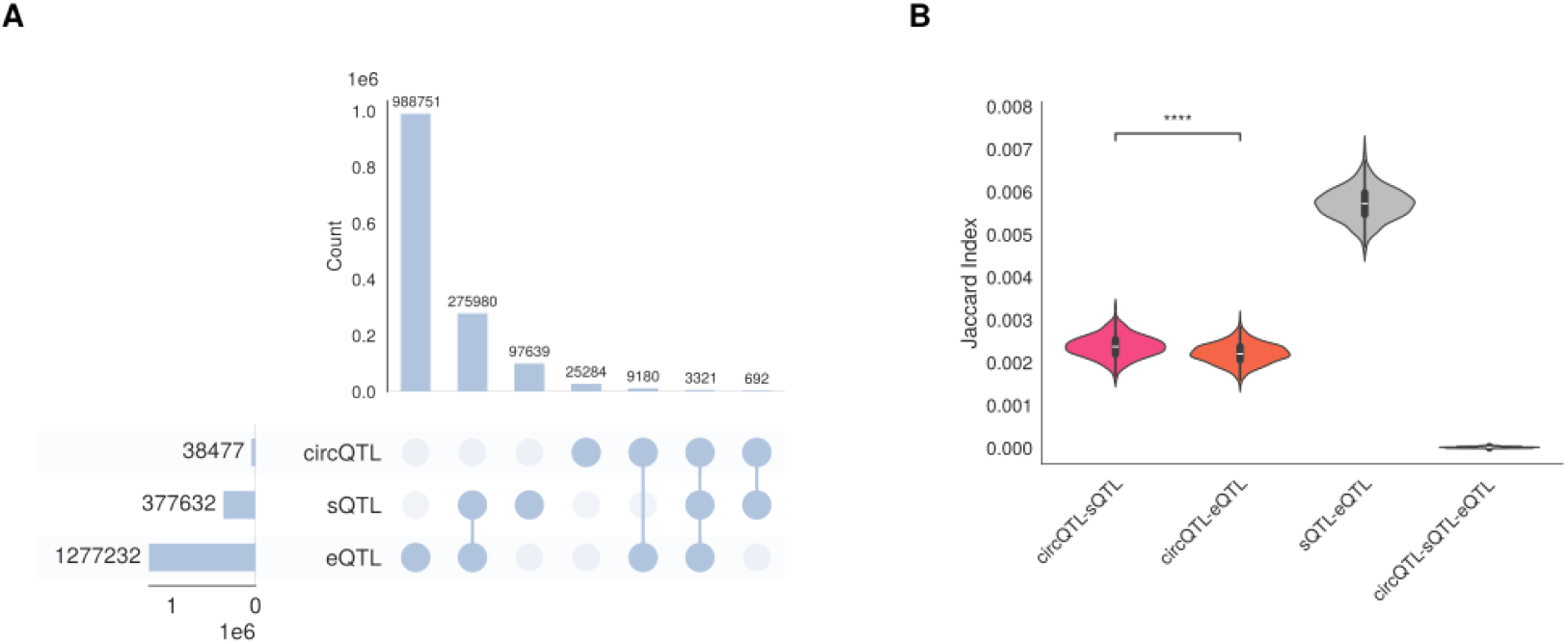
circQTL overlapping to sQTLs and eQTLs. (**A**) UpSetR visualization showing the overlap of circQTLs identified in our dataset with sQTLs and eQTLs, including pairwise and three-way intersections. Each vertical bar represents the size of a specific intersection, and the connected dots below indicate which QTLs are involved. Horizontal bars on the left indicate the total number of QTLs per group. (**B**) Violin plots showing bootstrap-derived Jaccard index distributions for pairwise and three-way comparisons among circQTLs, sQTLs, and eQTLs. Statistical significance between distributions was assessed using the Mann-Whitney U test; ****: *p ≤* 1.0 *×* 10^−4^.

### Genomic annotation reveals a circQTL enrichment in intergenic regions and a depletion in exon regions

The genomic location of a variant can influence its regulatory potential, affect splicing, or modulate gene expression. To determine whether circQTLs, sQTLs, and eQTLs follow distinct genomic patterns, we analyzed the distribution of their variant positions across gene element categories, including *introns, intergenic regions, exons, 3’-UTR, 5’-UTR, start codon*, and *stop codon*.

To focus on group-specific differences, we constructed a contingency table of absolute frequencies across gene element categories and applied a chi-square test of independence. This test revealed highly significant differences among the groups (*χ*^2^ = 2536.935, *df* = 12, *p <* 1 *×* 10^−300^) (**Figure 2A** and **Table S3**).

**Figure 2.**
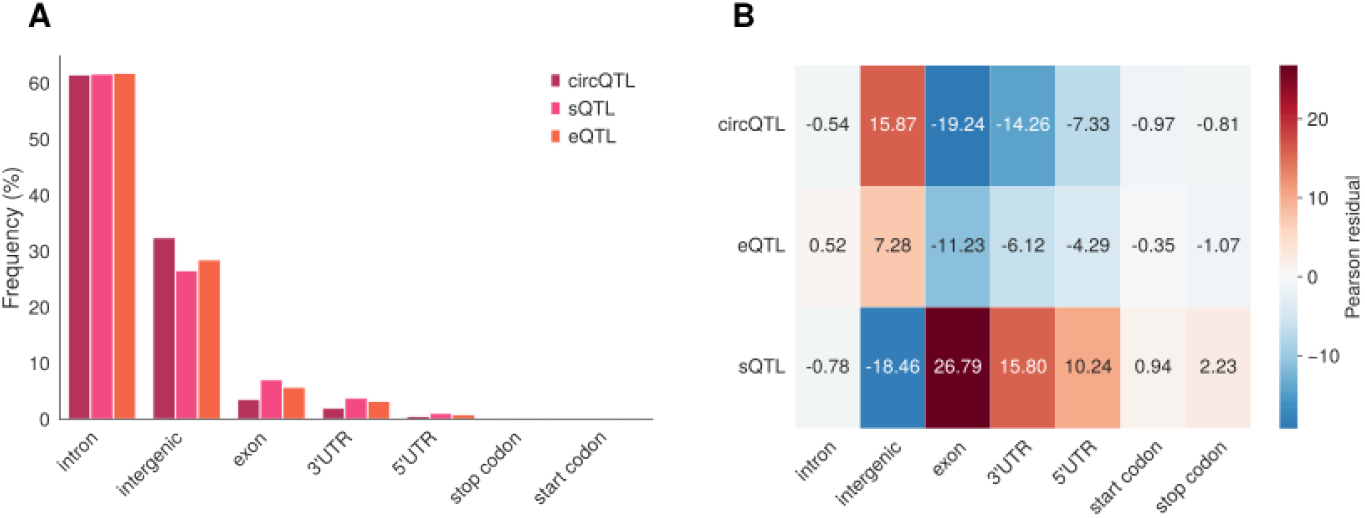
Genomic element annotation of circQTLs, sQTLs, and eQTLs. (**A**) Bar plot visualization showing the genomic element annotation frequency among circQTLs, sQTLs, eQTLs. (**B**) Heatmap of Pearson residuals showing enrichment and depletion of genomic elements across circQTLs, sQTLs, eQTLs. Residuals were calculated from a chi-square test of independence comparing the observed and expected frequencies of each gene element (columns) in circQTLs, sQTLs, and eQTLs (rows). Positive residuals (red) indicate enrichment, and negative residuals (blue) indicate depletion.

We then quantified the contribution of each genomic element to the overall chi-square statistic by calculating Pearson residuals. Positive residuals indicate enrichment relative to expectation, whereas negative residuals indicate depletion, allowing us to pinpoint which regions drive the differences among circQTLs, sQTLs, and eQTLs. Residuals showed that circQTLs were strongly enriched in intergenic regions (32.4%; *R* = 15.87) and depleted in exonic regions (3.6%, *R* = −19.24) relative to sQTLs and eQTLs. In contrast, sQTLs were enriched in exonic regions (7.0%, *R* = 26.79), whereas eQTLs exhibited a more balanced distribution across gene elements (exon *R* = −11.23, intron *R* = 7.28) (**Figure 2B**).

To identify which pairs of groups contributed most to these differences, we performed pairwise post-hoc chi-square tests with FDR correction. These analyses confirmed highly significant differences between circQTLs and sQTLs (*χ*^2^ = 1283.118, *FDR <* 1 *×* 10^−300^), circQTLs and eQTLs (*χ*^2^ = 731.945, *FDR* = 7.73 *×* 10^−155^), and sQTLs and eQTLs (*χ*^2^ = 1625.360, *FDR <* 1 *×* 10^−300^). Collectively, these results suggest that circQTLs preferentially localize to intergenic regions while avoiding exons, distinguishing them from sQTLs and eQTLs.

### Strong variation in circQTL overlap across validation datasets

To assess the reproducibility of circQTLs, we compared the circQTLs identified in our dataset with those reported in three independent validation datasets (Ahmed, Liu, and Mai). Overall, there is limited overlap across datasets, with only a small fraction of circQTLs shared with our dataset, specifically 6.55% for Liu et al., 2.34% for Ahmed et al., and 0.86% Mai et al. (**Figure 3A**). This observation indicates a strong dataset-specific bias in circQTL detection, likely reflecting differences in sample composition, sequencing depth, and detection pipelines.

**Figure 3.**
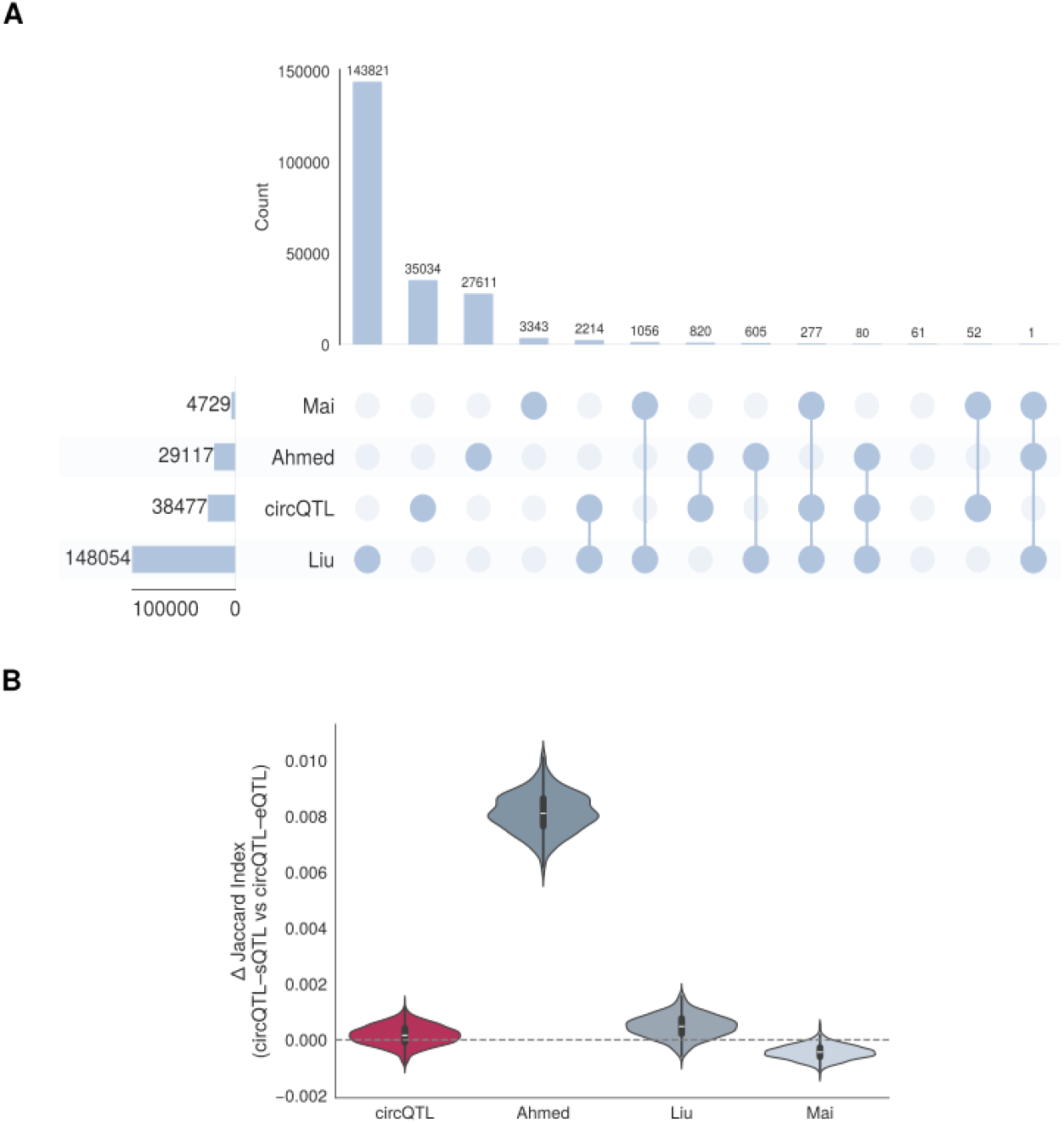
Validation of circQTLs across independent datasets. (**A**) UpSet plot showing the overlap of circQTLs identified in our dataset (circQTL) with three independent validation datasets (Ahmed, Liu, and Mai). Vertical bars represent the size of each intersection between datasets, while connected dots below indicate which datasets are included in the intersection. Horizontal bars on the left show the total number of circQTLs in each dataset. (**B**) Violin plot showing the bootstrap-derived Jaccard index differences between circQTL-sQTL and circQTL-eQTL comparisons across datasets. Positive values indicate stronger overlap with sQTLs, whereas negative values indicate stronger overlap with eQTLs.

In our dataset, the circQTL-sQTL Jaccard index is higher than for circQTL-eQTL (Δ*median* = 1.640 *×* 10^−4^), indicating stronger overlap with splicing QTLs. However, we wanted to demonstrate that relation in other datasets to validate whether this is characteristic of the biogenesis of circQTLs rather than a property of our dataset. The Ahmed (Δ*median* = 8.112 *×* 10^−3^) and Liu (Δ*median* = 4.830 *×* 10^−4^) dataset show a substantially higher circQTL-sQTL Jaccard index, whereas Mai (Δ*median* = −4.410 *×* 10^−4^) datasets show slightly negative differences (**Figure 3B, Figure S2** and **Table S4**).

Overall, these results support the general tendency of circQTLs to overlap more with sQTLs than with eQTLs, while also highlighting variability across independent datasets, which may reflect differences in sample size, experimental design, or tissue context.

## Discussion

circQTLs represent an emerging class of regulatory variants whose mechanistic basis has remained poorly defined. In this study, we systematically compared circQTLs with established sQTL and eQTL maps, showing that circQTLs exhibit a statistically significant but modestly stronger genomic overlap with sQTLs than with eQTLs (Mann–Whitney U test: *r* = 0.301). This pattern persists across independent validation datasets despite limited reproducibility of individual circQTLs between studies, suggesting that circRNA expression is preferentially influenced by splicing-associated mechanisms rather than transcriptional control of host genes. However, the modest effect size indicates that this preference is subtle and should be interpreted with caution.

CircRNA formation relies on back-splicing, a non-canonical splicing reaction in which a downstream splice donor is ligated to an upstream splice acceptor. This process directly competes with canonical pre-mRNA splicing for the same GT-AG splice sites and spliceosomal machinery (Chen and Yang, 2015; Zhang et al., 2016; Liang et al., 2017). For instance, Zhang et al. (2016) demonstrated that circRNA-producing genes exhibit faster Pol II elongation rates that provide critical temporal windows for intronic base-pairing and spatial juxtaposition of back-splice sites. In addition, Liang et al. (2017) found that the inhibition or slowing of co-transcriptional processing events shifts the transcript production from linear to circular isoforms.

In this scenario, variants affecting splice-site strength, intronic architecture, or spliceosome recruitment are therefore expected to influence the balance between linear and circular RNA production: a variant enhancing the effective strength or accessibility of splice sites in a back-splicing configuration favors circularization over linear splicing, while a weak allele may impair canonical splicing more strongly, potentially shifting the balance toward circularization under permissive kinetic or structural contexts. Our finding that circQTLs preferentially colocalize with sQTLs is consistent with a mechanistic coupling between circRNA biogenesis and splicing regulation, and suggests that splicing-associated regulatory variation may play a major role in shaping circRNA expression. However, additional functional and experimental validation will be required to establish the extent to which these effects are causal and to disentangle them from transcriptional contributions of host gene regulation.

Importantly, while circQTLs show a modest enrichment of colocalization with sQTLs relative to eQTLs, this does not imply that circQTLs are simply a subset of conventional sQTLs. Rather, a proportion of circQTLs likely reflect shared splice-site modulation affecting both linear splicing outcomes and back-splicing efficiency. However, our data also support the existence of circQTLs that do not exhibit detectable effects on canonical splicing metrics or gene expression, suggesting additional layers of regulatory specificity. These circQTLs may act through backsplicing-specific mechanisms that remain invisible to standard sQTL analyses, which are typically optimized for exon inclusion or linear junction usage and may not capture regulatory variation acting specifically on BSJ formation. Such mechanisms include modulation of intronic base-pairing, alteration of RNA secondary structure, recruitment of circRNA-promoting RBP, or changes in co-transcriptional kinetics that selectively impact the efficiency of circularization without perturbing linear isoform production.

This interpretation is further supported by the distinct genomic distribution of circQTLs. Compared to sQTLs, which show a clear enrichment in exonic regions consistent with direct splice-site modulation, we observed that circQTLs are strongly enriched in intergenic regions and depleted in exons. This pattern suggests that circRNA regulation frequently involves distal or structural regulatory elements rather than local exonic variation. Intergenic circQTLs may influence chromatin architecture, long-range enhancer-promoter interactions, or the spatial organization of intronic regions that facilitate back-splice site juxtaposition. Such regulatory modes align with accumulating evidence that circRNA biogenesis is sensitive to three-dimensional genome organization and intronic pairing rather than solely to local splice-site strength. Our observation of intergenic enrichment partially aligns with Ahmed et al. (2019). However, they reported enrichment across most genomic features except introns, a pattern not observed in our study

The limited overlap of circQTLs across independent datasets highlights substantial technical variability in detection, likely stemming from differences in sequencing depth, sample composition, and computational pipelines. Despite this, the global tendency of circQTLs to overlap more strongly with sQTLs than with eQTLs persists in Ahmed et al. (2019) and Liu et al. (2019) datasets, validating that enhanced splicing association represents a robust feature of circRNA genetic regulation rather than an artifact of our study design. The negative difference observed in the smaller trans-acting circQTLs dataset of Mai et al. (2022) highlights that distinct circQTL subclasses may exist, including variants whose downstream effects are mediated through circRNA-dependent regulation of distal genes rather than local splicing mechanisms.

Several limitations must be considered when interpreting these findings. First, circQTL detection remains highly constrained to accuracy and sensitivity of circRNA detection algorithms, sequencing depth, and back-splicing detection thresholds, which can substantially affect downstream association results as we see here. Benchmarking studies have consistently shown that different circRNA detection algorithms yield partially overlapping and often differ substantially in circRNA detection. Consensus-based protocols that integrate multiple callers can improve sensitivity and precision of circRNA detection and reduce tool-specific variability, suggesting that future circQTL studies may benefit from standardized multi-tool pipelines rather than relying on a single caller (Hansen, 2018; Vromman et al., 2023; Li et al., 2024; Zabala et al., 2026a). Second, conventional sQTL frameworks may underestimate the contribution of backsplicing-specific regulatory variants that do not perturb linear splicing patterns. Third, genomic annotation of intergenic circQTLs is inherently limited by incomplete knowledge of tissue-specific regulatory landscapes and chromatin interactions. Finally, the datasets used for external validation originate from heterogeneous biological contexts, including different tissues such as blood, brain, and lymphoblastoid cell lines (LCLs), and some cohorts contain samples from individuals with specific diseases (e.g., multiple sclerosis (MS) or autism). These differences in tissue type and disease status may introduce biological variability that complicates direct comparisons across studies and may influence the reproducibility of circQTL signals.

Some of these limitations may be resolved by integrating circQTLs with multi-omics data including chromatin accessibility (ATAC-seq), 3D genome architecture (Hi-C), and RBP binding profiles (eCLIP) to dissect distal regulatory mechanisms driving intergenic circQTL effects. Single-nucleus RNA sequencing (snRNA-seq) across tissues could reveal cell-type specific circRNA regulation patterns missed in bulk analyses. Experimental validation through CRISPR-based variant editing or splice-site perturbations in relevant cellular models will be essential to confirm causal links between identified circQTLs, splicing alterations, and circRNA biogenesis. In parallel, the development and adoption of standardized circQTL pipelines that combine multiple high-accuracy circRNA callers and re-quantification procedures are likely to reduce technical variability and improve cross-study comparability. Such integrative frameworks, similar in spirit to recently proposed consensus-based circQTL pipelines, could provide more coherent circQTL signals across tools and datasets and thereby strengthen downstream biological interpretation. Ultimately, incorporating circQTLs into disease GWAS colocalization frameworks may uncover novel regulatory axes contributing to complex trait heritability.

In conclusion, our results establish that circQTLs are preferentially associated with splicing-related regulatory variation rather than transcriptional control, positioning circRNA biogenesis firmly within the post-transcriptional regulatory layer of gene expression. At the same time, the existence of circQTLs without detectable linear splicing or expression effects suggests that circRNA regulation encompasses both shared and backsplicing-specific genetic mechanisms. Recognizing this dual architecture positions circQTLs as a distinct layer of genetic regulation operating at the interface between splicing kinetics, RNA structure, and genome organization.

## Methods

### CircRNA detection and quantification in RNA-seq data

We performed circRNA detection and quantification from peripheral blood leukocyte RNA sequencing (RNA-seq) data using data previously generated by our group and described in Iparraguirre et al. (2020). Briefly, we collected whole blood samples from people with MS and healthy controls (HC) at the Department of Neurology of Donostia University Hospital. In this study we analyzed the RNA-seq discovery cohort, which included 30 pwMS and 20 HC. We mapped RNA-seq reads to the hg19 genome assembly using BWA or Bowtie. We identified CircRNAs using `find_circ` (v1.0) (Memczak et al., 2013) and `CIRI2` (Gao et al., 2018), applying stringent filtering criteria requiring detection by both tools. We quantified circRNA expression using back-splice junction (BSJ) spanning reads obtained from CIRI2.

### circQTL calling

We conducted circQTL analysis by correlating the genotypes of 4,824,059 imputed autosomal variants with the expression levels of 22,308 circRNAs using the `MatrixEQTL` package (Shabalin, 2012). We restricted the analysis to cis-QTLs located within 1 Mb of the circRNA locus. We applied a linear regression model and controlled for group (MS and HC) and sex. We identified significant associations using a false discovery rate (FDR) threshold of < 0.05, accounting for the size of the discovery cohort, and we further filtered results by requiring a minor allele frequency (MAF) > 0.1. When multiple single-nucleotide polymorphism (SNP) candidates originated from the same genomic region, we calculated linkage disequilibrium (LD) using `LDmatrix`, focusing on the European population, and we retained SNPs with *r*^2^ *<* 0.7 to ensure independent inheritance. We originally aligned the dataset to the hg19 genome assembly and converted all genomic coordinates to the hg38 assembly using the UCSC `liftOver` tool (Rosenbloom et al., 2015).

### Comparison of circQTLs with sQTLs and eQTLs

We compared the identified circQTLs with sQTLs and eQTLs from the GTEx database to assess overlap and visualized the results using UpSet plots (Lex et al., 2014). To account for differences in sample size, we applied a bootstrap resampling procedure with *n* = 20,000 samples per iteration, based on an empirical evaluation of Jaccard index stability across increasing sample sizes (**Figure S1**). We repeated the analysis across 1,000 bootstrap iterations.

We calculated the Jaccard Index, which quantifies the overlap between two sets *A* and *B*, as

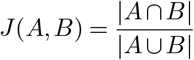

and, for comparisons involving three sets (*A, B*, and *C*), we computed the generalized Jaccard Index

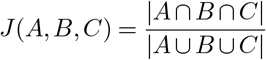

During each bootstrap iteration, we sampled elements with replacement from each set and calculated the Jaccard indices for the resampled data. This procedure generated bootstrap distributions for each pairwise and three-way comparison between circQTLs, sQTLs, and eQTLs. We assessed statistical differences between bootstrap distributions using the Mann-Whitney U test. In addition to statistical significance, we quantified the magnitude of the observed differences using the effect size (*r*) calculated as

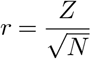

where *Z* is the standardized normal deviate derived from the Mann-Whitney U statistic, and *N* is the total number of observations across both groups (Fritz et al., 2012; Tomczak and Tomczak, 2014). This effect size provides a scale-independent measure of the strength of the difference between distributions. Bootstrap distributions and associated statistical results were visualized using violin plots.

### Genomic element annotation

We performed genomic element annotation using the GTF file corresponding to the GRCh38 (NCBI) version of the human genome. We systematically annotated variant positions according to genomic features, including *intron, intergenic, exon, 3’-UTR* (*3’ untranslated region*), *5’-UTR* (*5’ untranslated region*), *start codon* and *stop codon*.

Initially, all coordinates were classified as *intergenic*. Subsequently, coordinates were annotated based on detailed genomic features, including *3’-UTR, 5’-UTR, start codon* and *stop codon*. Positions not annotated in this step were then evaluated for overlap with *exon* regions. Those that still remained unannotated were next checked against *intronic* regions. Finally, any coordinates not matching any of the above categories were retained as *intergenic*. This stepwise annotation procedure was performed as in Zabala et al. (2026a).

We computed absolute frequencies and relative percentages for each genomic element across circQTLs, sQTLs, eQTLs, and we visualized these distributions using grouped bar plots.

To assess differences in genomic element composition among core QTL classes, we constructed contingency tables of absolute frequencies for circQTLs, sQTLs, and eQTLs. Given the discrete and count-based nature of the data, we evaluated overall distributional differences using the chi-square test of independence. We further quantified the contribution of individual genomic elements to the overall chi-square statistic using Pearson residuals, calculated for each cell of the contingency table as

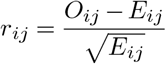

where *O*_*ij*_ is the observed count for group *i* and genomic element *j*, and *E*_*ij*_ is the expected count under the null hypothesis of independence. Positive values of *r*_*ij*_ indicate enrichment, while negative values indicate depletion of a given genomic element within a QTL group.

We performed post hoc pairwise chi-square tests between core QTL classes. We corrected all resulting p-values for multiple testing using the Benjamini-Hochberg false discovery rate (FDR) procedure.

### External validation

For external validation we used three datasets: the first, from 1000 Genomes lymphoblastoid cell lines, with 29117 circQTLs affecting circRNA expression independently of mRNA (Ahmed et al., 2019); the second, from 589 human dorsolateral prefrontal cortex samples, containing 148054 circQTLs and revealing technical, biological, and genetic factors influencing circRNA variation (Liu et al., 2019); and, the third, integrating RNA-seq and genotype data from autistic brains, including 4729 circQTLs involved in trans-eGene regulation and ASD-related causal pathways (Mai et al., 2022).

The dataset from Liu et al. (2019) was originally aligned to the hg19 genome assembly, and we converted all coordinates to the hg38 assembly using the UCSC liftOver tool.

## Abbreviations

BSJ: back-spliced junction
circRNA: circular RNA
circQTL: circular RNA-associated quantitative trait locus
eQTLs: expression quantitative trait locus
FDR: false discovery rate
GWAS: Genome-wide association studies
GTEX: Genotype-Tissue Expression Portal
HC: healthy control
LCL: lymphoblastoid cell line
LD: linkage disequilibrium
MAF: minor allele frequency
miRNA: microRNA
mRNA: messenger RNA
pre-mRNA: precursor messenger RNA
MS: multiple sclerosis
RBP: RNA-binding protein
RNA-seq: RNA sequencing
snRNA-seq: single-necleus RNA sequencing
QTL: quantitative trait locus
SNP: single-nucleotide polymorphism
sQTL: splicing quantitative trait locus
UTR: untranslated region

## Data availability

All data generated in this study are publicly available at Zenodo (Zabala et al., 2026b). The genomic annotation files used in this study are available at Zenodo Zabala et al. (2025). Whole-blood sQTL and eQTL datasets used in the analyses were obtained from the GTEx Portal (V10 release; QTL data downloads) on 4 December 2025.

## Code availability

All the code is available at Zenodo (Zabala et al., 2026c) and https://github.com/ZabalaAitor/circQTL-analysis.

## Author contributions

Author contributions are present in Table 1.

**Table 1.**
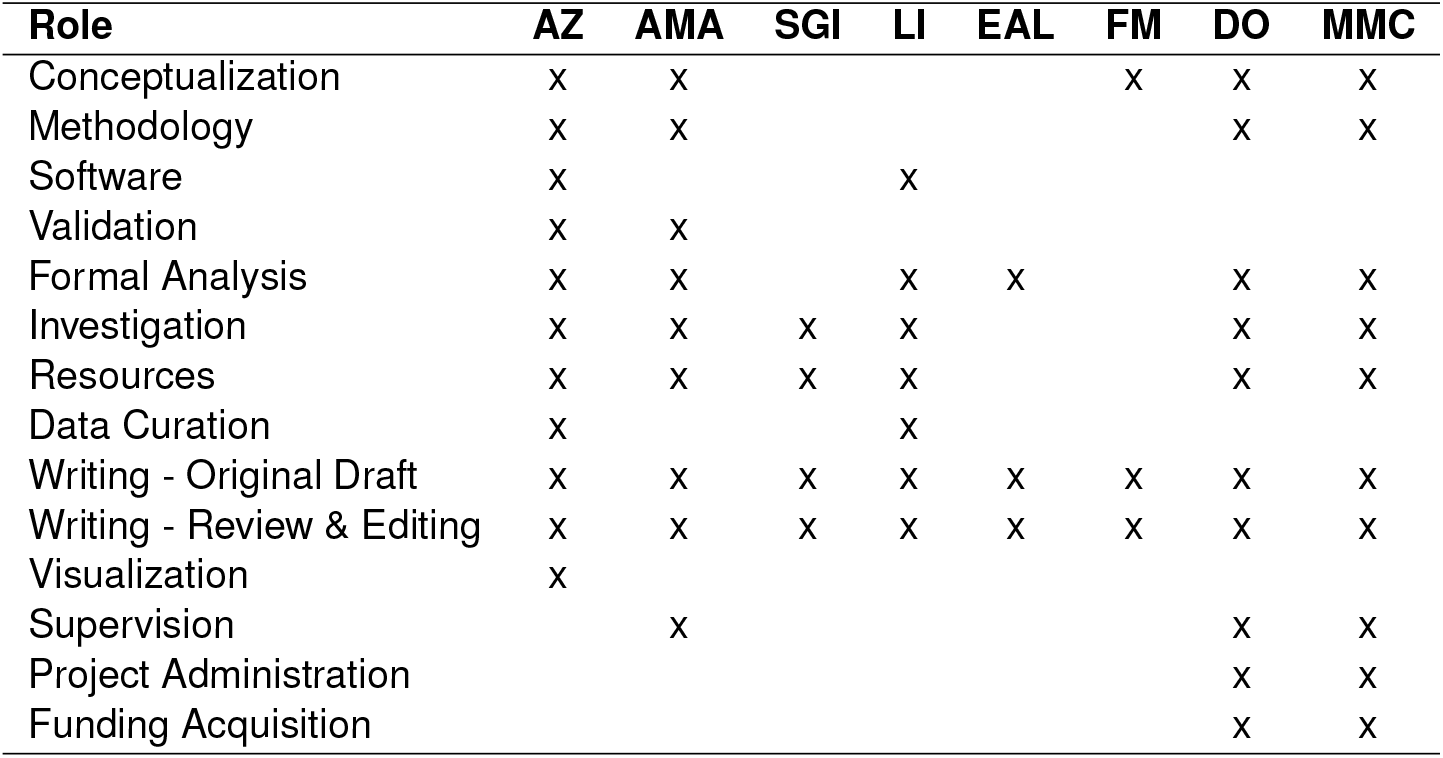
CRediT author contributions. An **X** indicates the involvement of each author in the corresponding contribution role. Descriptions of each role are provided below the table.

## CRediT role definitions

*Conceptualization*: Ideas; formulation or evolution of overarching research goals and aims.

*Methodology* : Development or design of methodology; creation of models.

*Software*: Programming and software development, algorithm design, code testing.

*Validation*: Verification of reproducibility or results.

*Formal Analysis*: Application of formal techniques to analyze or synthesize study data.

*Investigation*: Performing experiments or data/evidence collection.

*Resources*: Provision of study materials, instrumentation, or other tools.

*Data Curation*: Management activities to annotate, scrub data and maintain research data.

*Writing - Original Draft* : Writing the initial draft of the manuscript.

*Writing - Review & Editing*: Critical review and revision of the manuscript.

*Visualization*: Preparation or presentation of data visualizations.

*Supervision*: Oversight and leadership of the research activity.

*Project Administration*: Coordination responsibility for the research activity planning and execution.

*Funding Acquisition*: Acquisition of financial support for the project.

## Acknowledgments

The authors have used generative AI technology (ChatGPT) to improve the readability and overall quality of the manuscript. The use of AI has not altered the content or messages conveyed in the manuscript, focusing solely on refining the writing for clarity and enhanced readability.

## Supplementary Information

**Figure S1.**
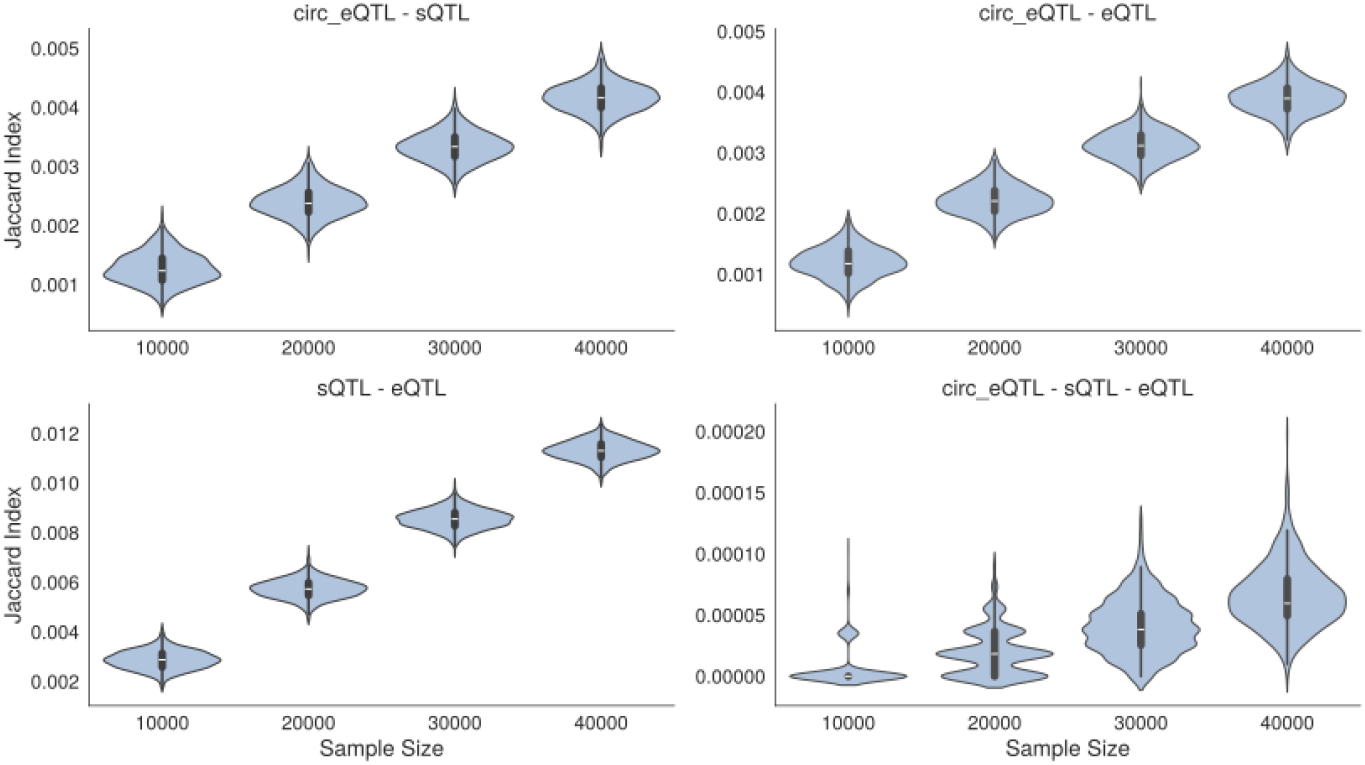
Bootstrap sample size evaluation for Jaccard overlap analysis.. Violin plots showing bootstrap-derived Jaccard index distributions across increasing sample sizes for pairwise and three-way comparisons among circQTLs, sQTLs, and eQTLs.

**Figure S2.**
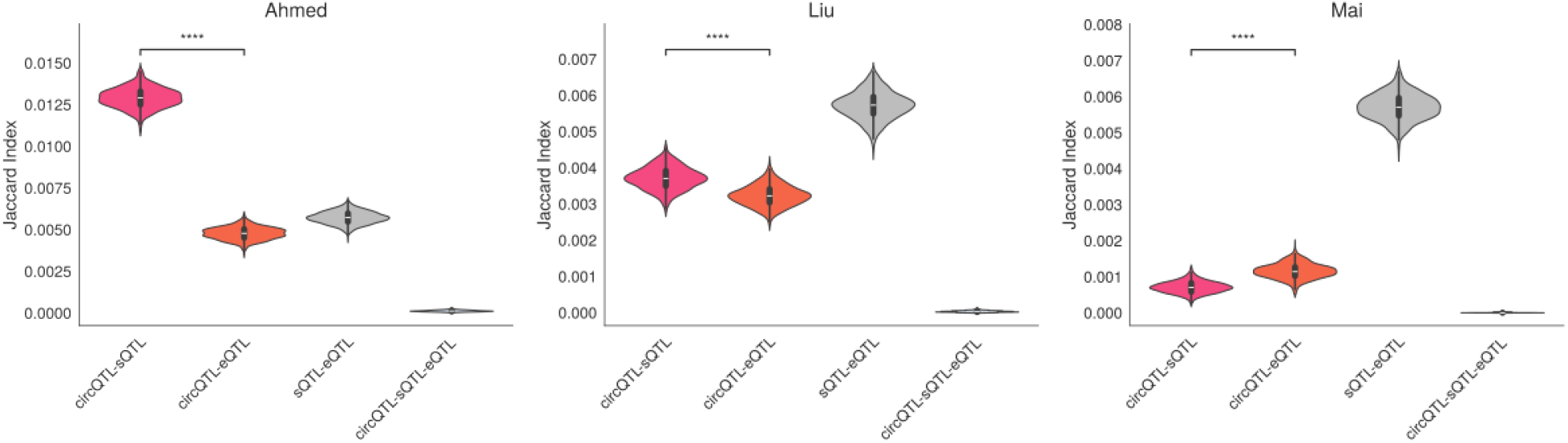
Validation of circQTL overlap with sQTLs and eQTLs across independent datasets. Violin plots showing bootstrap-derived Jaccard index distributions for pairwise and three-way overlaps among circQTLs, sQTLs, and eQTLs across validation datasets. Statistical significance between distributions was assessed using the Mann–Whitney U test; ****: *p* ≤ 1.0 *×* 10^−4^.

### Supplementary Tables

**Table S1.** Descriptive statistics of shared QTL overlaps. Summary of bootstrap-derived Jaccard index values for pairwise and three-way overlaps among circQTLs, sQTLs, and eQTLs across 1000 iterations.

| Statistic | circQTL–sQTL | circQTL–eQTL | sQTL–eQTL | circQTL–sQTL–eQTL |
| --- | --- | --- | --- | --- |
| Count | 1000 | 1000 | 1000 | 1000 |
| Mean | 0.002380 | 0.002213 | 0.005721 | 0.000021 |
| Std. Dev. | 0.000267 | 0.000243 | 0.000383 | 0.000019 |
| Minimum | 0.001571 | 0.001440 | 0.004573 | 0.000000 |
| 25% Quantile | 0.002201 | 0.002061 | 0.005472 | 0.000000 |
| Median | 0.002373 | 0.002207 | 0.005729 | 0.000018 |
| 75% Quantile | 0.002546 | 0.002377 | 0.005981 | 0.000037 |
| Maximum | 0.003348 | 0.003087 | 0.007116 | 0.000092 |

**Table S2.** Pairwise Mann-Whitney U test results for the Jaccard index. Two-sided non-parametric comparisons between the Jaccard index distributions of circQTL, sQTL, and eQTL overlaps. U statistic, P-value, and effect-size (*r*) are reported to indicate statistical significance and effect size for each pairwise comparison.

| Group 1 | Group 2 | U statistic | P-value | Effect size ($r$) |
| --- | --- | --- | --- | --- |
| circQTL–sQTL | circQTL–eQTL | 673653 | $3.18 \times 10^{-41}$ | 0.30 |
| circQTL–sQTL | sQTL–eQTL | 0 | $< 10^{-300}$ | -0.87 |
| circQTL–sQTL | circQTL–sQTL–eQTL | 1000000 | $< 10^{-300}$ | 0.87 |
| circQTL–eQTL | sQTL–eQTL | 0 | $< 10^{-300}$ | -0.87 |
| circQTL–eQTL | circQTL–sQTL–eQTL | 1000000 | $< 10^{-300}$ | 0.87 |
| sQTL–eQTL | circQTL–sQTL–eQTL | 1000000 | $< 10^{-300}$ | 0.87 |

**Table S3.**
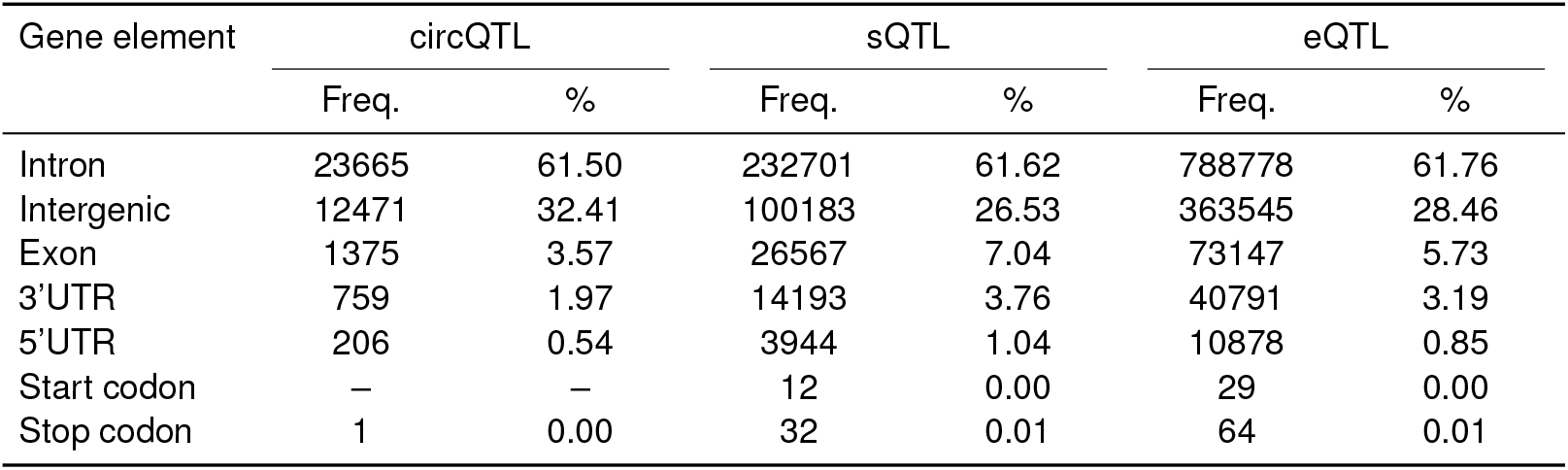
Genomic element annotation of circQTLs, sQTLs, and eQTLs. Distribution of variants across gene elements (intron, intergenic, exon, UTRs, start/stop codons) for each QTL type. Frequencies and percentages are relative to the total number of variants in each category.

| Gene element | circQTL |  | sQTL |  | eQTL |  |
| --- | --- | --- | --- | --- | --- | --- |
|  | Freq. | % | Freq. | % | Freq. | % |
| Intron | 23665 | 61.50 | 232701 | 61.62 | 788778 | 61.76 |
| Intergenic | 12471 | 32.41 | 100183 | 26.53 | 363545 | 28.46 |
| Exon | 1375 | 3.57 | 26567 | 7.04 | 73147 | 5.73 |
| 3'UTR | 759 | 1.97 | 14193 | 3.76 | 40791 | 3.19 |
| 5'UTR | 206 | 0.54 | 3944 | 1.04 | 10878 | 0.85 |
| Start codon | – | – | 12 | 0.00 | 29 | 0.00 |
| Stop codon | 1 | 0.00 | 32 | 0.01 | 64 | 0.01 |

**Table S4.** Distribution of differences between circQTL-sQTL and circQTL-eQTL overlaps. Summary statistics of the differences in Jaccard index across 1000 bootstrap iterations for each dataset (circQTL, Ahmed, Liu, Mai).

| Dataset | Count | Mean | SD | Min | 25% | Median | 75% | Max |
| --- | --- | --- | --- | --- | --- | --- | --- | --- |
| circQTL | 1000 | 0.000167 | 0.000359 | -0.001001 | -0.000087 | 0.000164 | 0.000419 | 0.001387 |
| Ahmed | 1000 | 0.008128 | 0.000714 | 0.005860 | 0.007653 | 0.008112 | 0.008647 | 0.010338 |
| Liu | 1000 | 0.000483 | 0.000413 | -0.000831 | 0.000216 | 0.000483 | 0.000765 | 0.001800 |
| Mai | 1000 | -0.000449 | 0.000259 | -0.001340 | -0.000607 | -0.000441 | -0.000275 | 0.000589 |

